# Tasmanian devil CD28 and CTLA4 capture CD80 and CD86 from adjacent cells

**DOI:** 10.1101/2020.06.11.145789

**Authors:** Candida Wong, Jocelyn M. Darby, Peter R. Murphy, Terry L. Pinfold, Patrick R. Lennard, Gregory M Woods, A. Bruce Lyons, Andrew S. Flies

## Abstract

Immune checkpoint immunotherapy is a pillar of human oncology treatment with potential for non-human species. The first checkpoint immunotherapy approved for human cancers targeted the CTLA4 protein. CTLA4 can inhibit T cell activation by capturing and internalizing CD80 and CD86 from antigen presenting cells, a process called trans-endocytosis. Similarly, CD28 can capture CD80 and CD86 via trogocytosis and retain the captured ligands on the surface of the CD28-expressing cells. The wild Tasmanian devil (*Sarcophilus harrisii*) population has declined by 77% due to transmissible cancers that evade immune defenses despite genetic mismatches between the host and tumours. We used a live cell-based assay to demonstrate that devil CTLA4 and CD28 can capture CD80 and CD86. Mutation of evolutionarily conserved motifs in CTLA4 altered functional interactions with CD80 and CD86 in accordance with patterns observed in other species. These results suggest that checkpoint immunotherapies can be translated to evolutionarily divergent species.

**Highlights:**

- Key immune checkpoint receptor-ligand interactions are conserved in marsupials.
- Live cell-based assays show Tasmanian devil CD28 and CTLA4 can capture CD80 and CD86 *in trans* from adjacent cells.
- Mutation of the conserved CTLA4_MYPPPY_ ligand binding motif to CTLA4_MYPPPA_ reduces binding to CD80 and intercellular protein transfer.
- Removal of conserved CTLA4_YVKM_ protein recycling binding motif in CTLA4 results in bidirectional intercellular protein transfer between CTLA4 and CD80.
- Highly successful human immune checkpoint immunotherapies have the potential to be translated for veterinary and conservation medicine.

## 1. Introduction

The initial USA Food and Drug Administration approval for an immune checkpoint inhibitor was in 2011 when a monoclonal antibody targeting CTLA4 was approved for metastatic melanoma (U.S. Food and Drug Administration, 2011). Since then, targeting immune checkpoints has proved effective for many types of cancer, and a single monoclonal antibody targeting the PD1 pathway is currently approved for 15 types of cancer (U.S. Food and Drug Administration, 2020). Despite the massive potential for veterinary medicine and One Health projects, functional comparative studies of immune checkpoints studies are limited, and have focused primarily on domestic animals (Achleitner et al., 2011; Bernard et al., 2007; Folkl et al., 2010; Hansen et al., 2009; Hartley et al., 2016; Ikebuchi et al., 2014, 2013, 2011; Maekawa et al., 2014, 2017, 2016; Nemoto et al., 2018; Rebech et al., 2020; Tagawa et al., 2016).

The Tasmanian devil (*Sarcophilus harrisii*) immune system has received attention in recent years due to two transmissible cancers that have decimated the wild Tasmanian devil population (Pearse and Swift, 2006; Pye et al., 2016). The original transmissible devil facial tumor (DFT1) was first observed in 1996 and has since spread across most of the island state, leading to an average 77% population decline in areas were the DFT1 is present (Lazenby et al., 2018). In 2014, a second independent transmissible devil facial tumor (DFT2) was discovered in southern Tasmania (Pye et al., 2016). DFT1 and DFT2 are lethal and the available evidence suggests that both tumors have arisen from Schwann cells (Murchison et al., 2010; Patchett et al., 2020). A primary means of immune evasion by DFT1 cells is downregulation of MHC-I. However, MHC-I can be upregulated in response to IFNγ (Siddle et al., 2013). The DFT2 cells examined to date generally express MHC-I (Caldwell et al., 2018). This suggests that these transmissible cancers employ additional immune evasion mechanisms to induce tolerance to tumor allografts and damage-associated molecular patterns.

We have shown that the immune checkpoint protein PDL1 binds to PD1 on DFT cells and developed devil-specific monoclonal antibodies that can block this receptor-ligand interaction (Flies et al., 2020a, 2016). Furthermore, exposure to IFNγ upregulates PDL1 on DFT cells, matching the pattern observed in many human and mouse cancers. Comparative sequence analysis of other key checkpoint proteins in devils showed that key regions in mouse and human CD28 and CTLA4 proteins are conserved in devils (Flies et al., 2017). Additionally, binding of CTLA4 to its ligands CD80 and CD86 has been confirmed using soluble recombinant proteins (Flies et al., 2020a). However, functional analyses of cell surface protein interactions and protein trafficking remain unexplored.

In humans and mice, CTLA4 can capture CD80 and CD86 from the surface of adjacent cells via trans-endocytosis, which results in internalization and degradation of CD80 and CD86 in the capturing cell. CTLA4 is continuously recycled from the cell surface to endosomal compartments, which allows a single CTLA4 protein to capture multiple ligands (Qureshi et al., 2011; Robles et al., 2012). CTLA4-mediated depletion of CD80 and CD86 from antigen presenting cells results in reduced T cell activation due to impaired CD28 costimulation (Qureshi et al., 2011).

CD28 is known to capture CD80 and CD86 from the surface of adjacent cells, but the captured ligands remain bound to CD28 on the surface of the capturing cell (Briggs, 2014; Hwang et al., 2000; Qureshi et al., 2011). This process of trogocytosis has also been well-documented for capture of MHC-I and MHC-II by T cells, NK cells, and basophils, and can result in immune suppression due to depletion of MHC from target cells (Akkaya et al., 2019; Chung et al., 2014; Daubeuf et al., 2006; Gu et al., 2012; Joly and Hudrisier, 2003; Lemaoult et al., 2015; Nakayama et al., 2011).

We developed a panel of immune checkpoint proteins fused to fluorescent reporter proteins and used coculture assays to monitor protein transfer using flow cytometry to determine if intercellular protein transfer patterns evolutionarily conserved in Tasmanian devils. This approach revealed that CTLA4 and CD28 can capture costimulatory ligands from neighboring cells, and that key protein motifs for ligand binding and protein trafficking were conserved in devils. The conserved functional interactions of these proteins across the 160 million years since the divergence of eutherian and metatherian mammals (Luo et al., 2011) suggests strong stabilising selection and that there is potential to replicate successful human immunotherapies in devils and other species.

## 2. Materials and methods

### 2.1 Design of expression vectors

Complete methodological details with step-by-step instructions for (a) *in silico* characterization of protein structure, (b) design and (c) assembly of expression vectors, and (d) transfection of mammalian cells has been previously published (Flies et al., 2020a, Flies et al., under review at Bio-protocol).

Open reading frames for the devil genes of interest (CD28, CTLA4, CD80, and CD86) were retrieved from Genbank and Ensembl databases (Table S1). *In silico* analysis of the protein coding sequences for genes-of-interest in this study were performed in a previous study (Flies et al., 2017). The expression vector design included the full-length open reading frames for gene-of-interest fused to a tobacco etch virus cleavage tag, a linker peptide, a 6x-Histidine purification tag, and monomeric fluorescent reporter protein. The direct fusion of the protein-of-interest to a fluorescent reporter protein allow for protein tracking in live cells (Flies et al., 2020a). The vectors contain an all-in-one transfection system that includes antibiotic resistance and genes-of-interest within a Sleeping Beauty transposon cassette (Kowarz et al., 2015), and the Sleeping Beauty transposase outside of the transposon cassette (Mátés et al., 2009). CTLA4, CD28, CD80, and CD86 were fused to mCitrine (Addgene # 135923), mOrange (Addgene # 135928), mTagBFP (Addgene # 135924), and mNeptune2 (Addgene # 135927), respectively.

### 2.2 Restriction digest of expression vectors

2 μg of base expression vectors were digested using NotI-HF (NEB # R3189S) and SmaI (NEB # R0141S) overnight at room temperature. Antarctic phosphatase (NEB # M0289S) was then added to the vector digests and incubated for 60 minutes at 37°C, then for a further 2 minutes at 80°C to inactivate the enzymes. The reactions were then run at 100V in a 1% agarose gel and the digest vectors were excised and purified using a NucleoSpin^®^ Gel and PCR Clean-up Kit (Macherey-Nagel, USA) according to instructions set by the manufacturer. DNA concentrations were measured and analysed using a NanoDrop(tm) 1000 Spectrophotometer (NanoDrop Technologies, USA).

### 2.3 Overlap extension PCR to create vector inserts

Overlap extension PCR was used to extend open reading frames and remove the stop codon from the genes of interest to allow for Gibson assembly of plasmids pCW1, pCW2, pCW3, and pCW5 (Gibson et al., 2009). Primers and reaction conditions are available in Table S2. PCR amplicons were isolated using gel electrophoresis run at 100 V for 30 minutes in a 1% agarose gel. DNA was purified and quantified as described above. CTLA4 mutants with a truncated C-terminus (pCW9; CTLA4_truncated_; amino acids 211-223 deleted) or the YVKM protein trafficking motif deleted (pCW8; CTLA4_del_; amino acids 201-223 deleted) were developed as previously described by Qureshi et al. (2012) with minor modifications. pCW8 and pCW9 were made by amplifying with a reverse primer that amplified the DNA upstream of the deleted region and extension that overlapped the expression vector (Table S2). The CTLA4 expression vector (pCW10) with an altered binding motif was produced by substituting an alanine for the for the second tyrosine in the MYPPPY ligand binding motif (Y139A) to create the CTLA4_MYPPPA_ mutant (Morton et al., 1996). The mutation was coded into new forward and reverse primers (Table S2), and were used to amplify and extend one fragment upstream of the mutation, and one fragment downstream of the mutation. The two fragments were purified, combined in a PCR reaction, and re-amplified using the forward from the 5’ fragment and the reverse primer from the 3’ fragment to form one large fragment. This fragment was purified and assembled using NEBuilder as described above.

### 2.4 Assembly and transformation of expression vectors

The DNA for the gene-of-interest with overlap extensions were ligated into the digested vectors using NEBuilder® HiFi DNA Assembly Cloning Kit (NEB # E5520S) by incubating at 50 °C for one hour according to the manufacturer’s instructions. A negative control was established by transforming DH5α with the linearised vector. A positive control on the other hand was established by transforming DH5α bacteria with NEBuilder® Positive Control (New England Biolabs, USA). After transformation, the bacteria were grown overnight at 37°C on Luria Broth Agar plates added with 100 μg/mL of ampicillin.

Colony PCR was used to identify colonies with the predicted insert size. 10 μL of OneTaq Quick-Load 2X Master Mix (NEB # M0486L) was used for all colony PCR. 7 μL of deionized-distilled water was added to the mix, and then 1 μL of forward (pSB_EF1a.FOR: GCCTCAGACAGTGGTTCAAAG) and reverse (pSB_BGH.REV: AGGCACAGTCGAGGCTGAT) primers at 10 μM concentration were diluted into the mix. 4 to 6 bacterial colonies were picked using sterile pipette tips and transferred to separate tubes containing 10 µL of PBS for assessing. 1 uL of bacteria in PBS was then transferred to the PCR tube to achieve a final volume of 20 μL and primer concentration of 0.5 μM. The mix was then denatured at 94 °C for 5 minutes, and then subjected to 30 cycles of denaturation at 94 °C for 15 seconds, annealing at 58 °C for 15 seconds, and extension at 68 °C for 105 seconds, before a final extension step at 72 °C for 3 minutes. 10 μL of the reaction was then added directly to a 1% agarose gel and run at 100 V for 25-35 minutes. DNA bands were imaged under UV light using Gel Doc(tm) XR+ Gel Documentation System (Bio-Rad, USA).

### 2.5 Purification and sequencing of expression vectors

Colonies that yielded amplicons that matched the predicted amplicon size were cultured overnight for plasmid amplification. 9 µl of bacteria in PBS was added to 5 mL of Luria Broth with 100 μg/ml of ampicillin. The bacterial cultures were incubated at 37°C and 200 rpm overnight. The plasmids were isolated from bacterial culture using PureLink(tm) Quick Plasmid Miniprep Kit (Invitrogen, USA) or NucleoSpin® Plasmid EasyPure Miniprep Kit (Macherey-Nagel, USA) according to the manufacturers’ instructions. Concentrations of purified plasmids were quantified using a NanoDrop(tm) 1000 Spectrophotometer (NanoDrop Technologies, USA). The plasmids were stored at −20°C. The plasmids were sequenced using a BigDye(tm) Terminator v3.1 Cycle Sequencing Kit (Applied Biosystems, USA). The BigDye(tm) Terminator was removed using Agencourt® CleanSEQ® (Beckman Coulter, USA) according to the protocols set by the manufacturer before loading samples to the Applied Biosystems® 3500xL Genetic Analyzer (Applied Biosystems, USA) for sequencing using a fluorescence-based capillary electrophoresis analytical technique.

### 2.6 Transfection of Chinese Hamster Ovary (CHO) cells

CHO-K1 (ATCC CCL-61) cells were cultured in cRF10 (complete RPMI media supplemented with 10% fetal bovine serum) in 75 cm^2^ culture flasks (Corning, USA) at 37°C in a humidified atmosphere with 5% CO_2_. CHO cells were transfected using a modified polyethylenimine-based (PEI) method. 300,000 cells/well were seeded into 6-well plates and were grown overnight at 37°C with 5% CO_2_. 2 μg of plasmid DNA and 6 μg of PEI were diluted separately to 100 μL of PBS in separate microfuge tubes. The solution from the DNA tube was then added to PEI tubes and mixed by gentle pipetting. Tubes with a total volume of 200 µl of reagents were incubated for 15 to 20 minutes at room temperature. While the DNA:PEI solution was incubating the media on the target CHO cells in 6-well plates was replaced with fresh cRF10. The transfection was accomplished by adding the entire contents of DNA:PEI solution (200 µL) dropwise to the cells. Cells were then incubated with transfection reagents overnight and examined for fluorescence on the Leica-DM IRB fluorescence microscope (Leica, Germany). After fluorescence was detected indicating that transfection was successful, the media was removed and replaced with cRF10 containing 1 mg/mL of hygromycin B solution from *Streptomyces hydroscopicus* (Invitrogen, USA). Media was refreshed as needed until drug selection was complete and all cells were fluorescent. The cells were then cultured in cRF10 with a maintenance dose of 0.2 mg/mL of hygromycin B.

### 2.7 Intercellular protein transfer coculture assay

The coculture assay to assess intercellular protein transfer was modified from Briggs (2014). The lysosomal inhibitor chloroquine was used to block degradation of captured proteins that were endocytosed (Qureshi et al., 2011). Chloroquine was diluted to 100 µM in cRF5 (complete RPMI with 5% fetal bovine serum) and 100 µL were added into appropriate wells in a 96-well round bottom plate (Costar, USA). The plate was incubated at 37°C with 5% CO_2_ until cell lines were ready to be cocultured. Stably-transfected CHO cells were harvested, washed, and resuspended to 5 × 10^5^ cells/mL in cRF5. 50 µL (25,000 cells) of cell line A (e.g. CD28) and cell line B (e.g. CD80) were aliquoted into appropriate wells to achieve a 1:1 ratio and a total of 50,000 cells/well. The final volume was 200 µL/well with a final chloroquine concentration of 50 µM. The plates were then incubated at 37°C with 5% CO_2_ for either 3 hours or 20 hours.

### 2.8 Flow cytometry

The coculture assay plates were centrifuged at 500g for 3 minutes at 4°C. The 96-well plate was then inverted, and the media was discarded by briskly dumping the media out. 100 µL of PBS was added into appropriate wells to wash cells and the plate was spun again at 500g for 3 minutes at 4°C. After removing media, 100 µL of Trypsin-EDTA was added into each well and incubated at 37°C with 5% CO_2_ for 5 to 10 minutes or until cells were dislodged from the bottom of the well. 150 µL of cRF10 was then added to each well and the cells were resuspended by pipetting. The plate was then centrifuged at 500g for 3 minutes at 4°C and media was discarded. 200 µL of flow cytometry buffer was added into each well. Following this, the plate was spun again at 500g for 3 minutes at 4°C. Media was discarded and cells were resuspended in 250 µL of flow cytometry fixation buffer (Flies et al., 2020a). The coculture plate was placed on a rocking platform and incubated at room temperature and protected from light for at least 20 minutes. The incubated plate was subsequently spun at 500g for 3 minutes at 4°C and the media was discarded (Flies et al., 2020a). Cells in each well were resuspended in 250 µl of flow cytometry buffer, wrapped in foil, and stored at 4°C for at least 2 hours before flow cytometric analysis.

### 2.9 Data Processing and Gating Strategy for Flow Cytometry Analysis

Flow cytometry data was analysed using the FCS Express 6 Flow Cytometry Software version 6 (De Novo Software). CHO cells were primarily gated by forward scatter height (FSC-H) and side scatter height (SSC-H) to exclude cell debris, and then sub-gated for singlets using FSC-A (area) and FSC-H. CHO cells that did not express a fluorescent protein or expressed a single fluorescent protein were used as controls for setting voltages and compensation. A minimum of 10,000 cells was acquired from each experimental sample, and all cell combinations were done with n=3 technical replicates.

Intercellular protein transfer was identified according to the schematic in **Fig. 1A**. The upper left quadrant (UL) and lower right quadrant (LR) indicate cells that have only a single color of fluorescent protein (**Fig. 1A**). The upper right quadrant (UR) indicates cells that are positive for two colors, indicating the intercellular transfer of a fluorescent protein occurred (Briggs, 2014). Additionally, a shift in both cell lines to the UR quadrant indicates a bidirectional protein exchange, whereas a shift from one of the cell lines to the UR quadrant indicates a one-way protein transfer (**Fig. 1A**).

**Figure 1.**
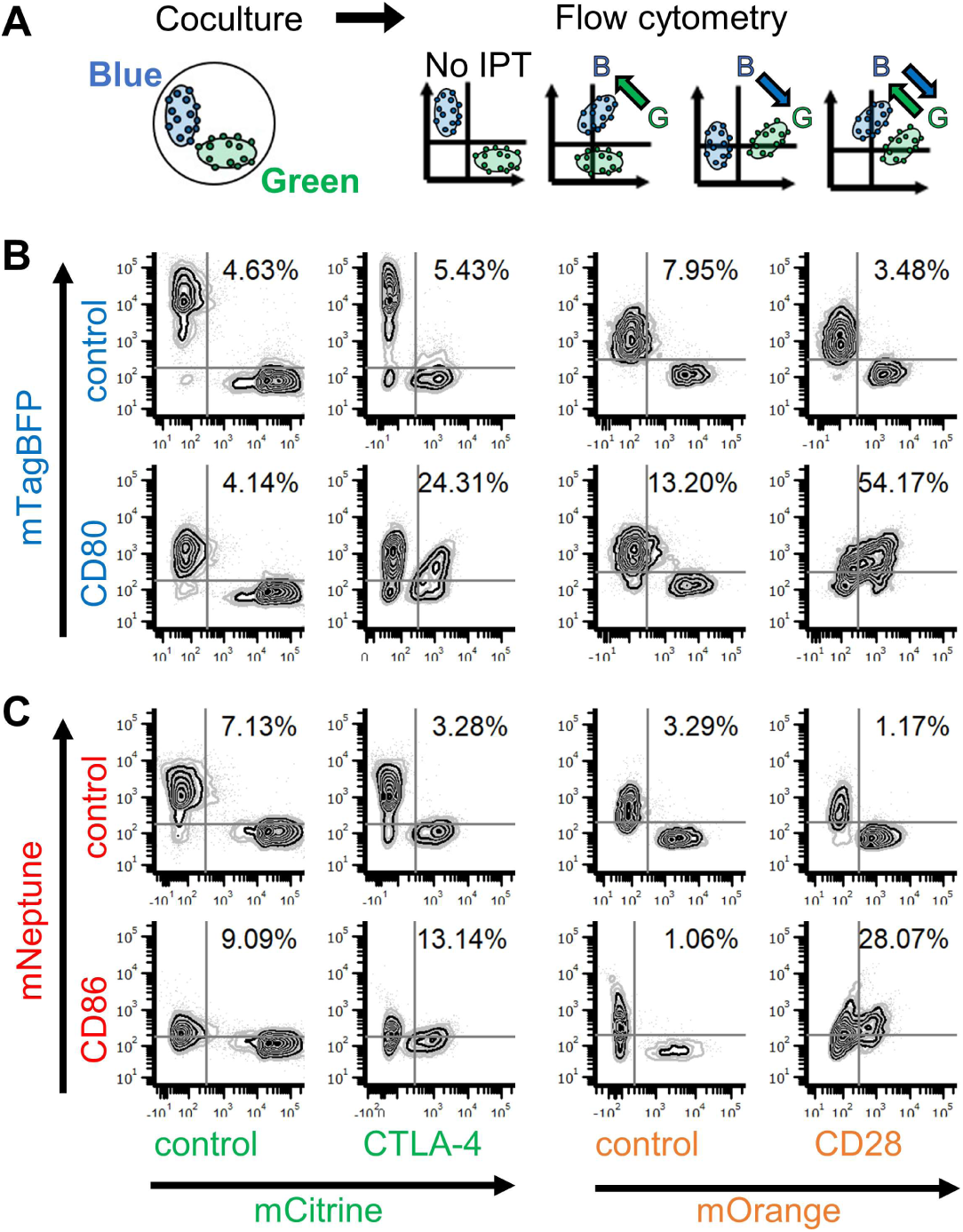
Schematic of intercellular protein transfer (IPT) and checkpoint interactions. (A) Cells expressing fluorescent fusion proteins (e.g. blue and green) are cocultured in a 96-well plate with chloroquine to block lysosomal degradation of internalized proteins. Flow cytometry is used to analyze IPT. No IPT is observed when cells remain in upper left (UL) and lower right (LR) quadrants (no shift). One-way protein exchange is observed if either blue or green cells shift to upper right (UR) quadrant. Bidirectional or two-way protein exchange is observed if both blue and green cells shift to UR quadrant. (B) Flow cytometry plots of CTLA4-mCitrine and CD28-mNeptune cocultured with CD80-mTagBFP; mTagBFP, mCitrine, and mOrange were used as controls in a 20-hour assay. IPT is observed between CTLA4 and CD80 (one-way), and CD28 and CD80 (bidirectional). Results shown are representative of n=2 technical replicates.

### 2.10 Analysis of CTLA4 binding and protein trafficking motifs across species

CTLA4 protein sequences were obtained from the National Center for Biotechnology Information (NCBI) (https://www.ncbi.nlm.nih.gov/gene/1493/ortholog/). Descriptive information for CTLA4 for each class (*Amphibia, Reptilia, Aves, Mammalia*) were exported separately from NCBI as tabular data (e.g. comma separated values spreadsheet). The files were manually updated to include a new column containing the class (e.g. *Aves*). The descriptive spreadsheets were then combined to create a spreadsheet with descriptive information for all species. The RefSeq protein sequences were exported from NCBI as a FASTA file. FASTA sequences were imported into CLC Main Workbench (CLC, 2020), and the “Show Table” tab was used to determine the length of each CTLA4 protein sequence. This table was copied and saved as a .CSV file for subsequent merging with other spreadsheets. The “Motif Search” function in CLC was then used to identify and annotate all sequences containing a greater than 90% match to the CTLA4_MYPPPY_ and CTLA4_YVKM_ motifs. A new .CSV file was created for the CTLA4_MYPPPY_ and the CTLA4_YVKM_ matches. The four .CSV files were then merged in R version 3.6.1 using the “merge” function (R Core Team, 2019). The script is available with the supplementary material (r_CTLA4_merge_master.R) and the merged data are available in Table S3.

## 3. Results and Discussion

CHO cells expressing either unfused mCitrine or mTagBFP controls resulted in the cells remaining primarily positive for only a single color after coculture (**Fig. 1B**). Likewise, cells remained mostly single color positive after coculture of CTLA4-mCitrine with mTagBFP control or mCitrine control with CD80-mTagBFP (**Fig. 1B**). However, coculture of CTLA4-mCitrine with CD80-mTagBFP for 20 hours resulted in 24% of the cells in the upper right quadrant. CTLA4 cells shifted from the lower right quadrant to the upper right quadrant indicating positive receptor-ligand binding and protein transfer from CD80-mTagBFP to CTLA4-mCitrine (**Fig. 1B**). The coculture between cells transfected with CD28-mOrange and CD80-mTagBFP suggests that bidirectional protein transfer occurred as both single color populations shifted, with 54% of the cells were in the upper right quadrant after coculture. CTLA4 also captured CD86, whereas transfer of CD28 and CD86 again appeared to be bidirectional (**Fig. 1C**). The interaction of CTLA4 and CD28 with CD80 appears stronger than with CD86, but this could be an artefact of the suboptimal excitation of the mNeptune fluorescent protein; mNeptune has peak excitation at 600 nm, but our available flow cytometer had a 633 nm laser.

To assess the role of key functional motifs in CTLA4 we developed three CTLA4 mutants (**Fig. 2A**). Here, we used 3-hour and 20-hour coculture assays to document IPT over time, and also to minimise potential side effects of extended cell culture with chloroquine. The results show a one-way IPT from CD80-mTagBFP to CTLA4-mCitrine in 3-hour (17%) and 20-hour (46%) assays (**Fig. 2B**). The devil mutant truncated CTLA4_truncated_-mCitrine (**Fig. 2B**) also showed one-way protein exchange in both 3-hour (16%) and 20-hour (44%) assays with CD80 (**Fig. 2B**). The YVKM motif in the cytoplasmic domain of CTLA4 is important for protein recycling, so we hypothesized that CTLA4-CD80 binding would remain intact in a CTLA4 mutant with the YVKM motif removed, but that CTLA4 would have reduced capacity to trans-endocytose CD80 and CD86. As expected, deletion of the YVKM dramatically resulted in a pattern more closely resembling the interaction of CD28-CD80 than the CD80 with the other CTLA4 variants (**Fig. 2B**). The CTLA4 and CD86 interaction was also changed in the CTLA4_YVKM-del_ mutant, with the CTLA4 and CD86 populations merging towards a single population after coculture.

**Figure 2.**
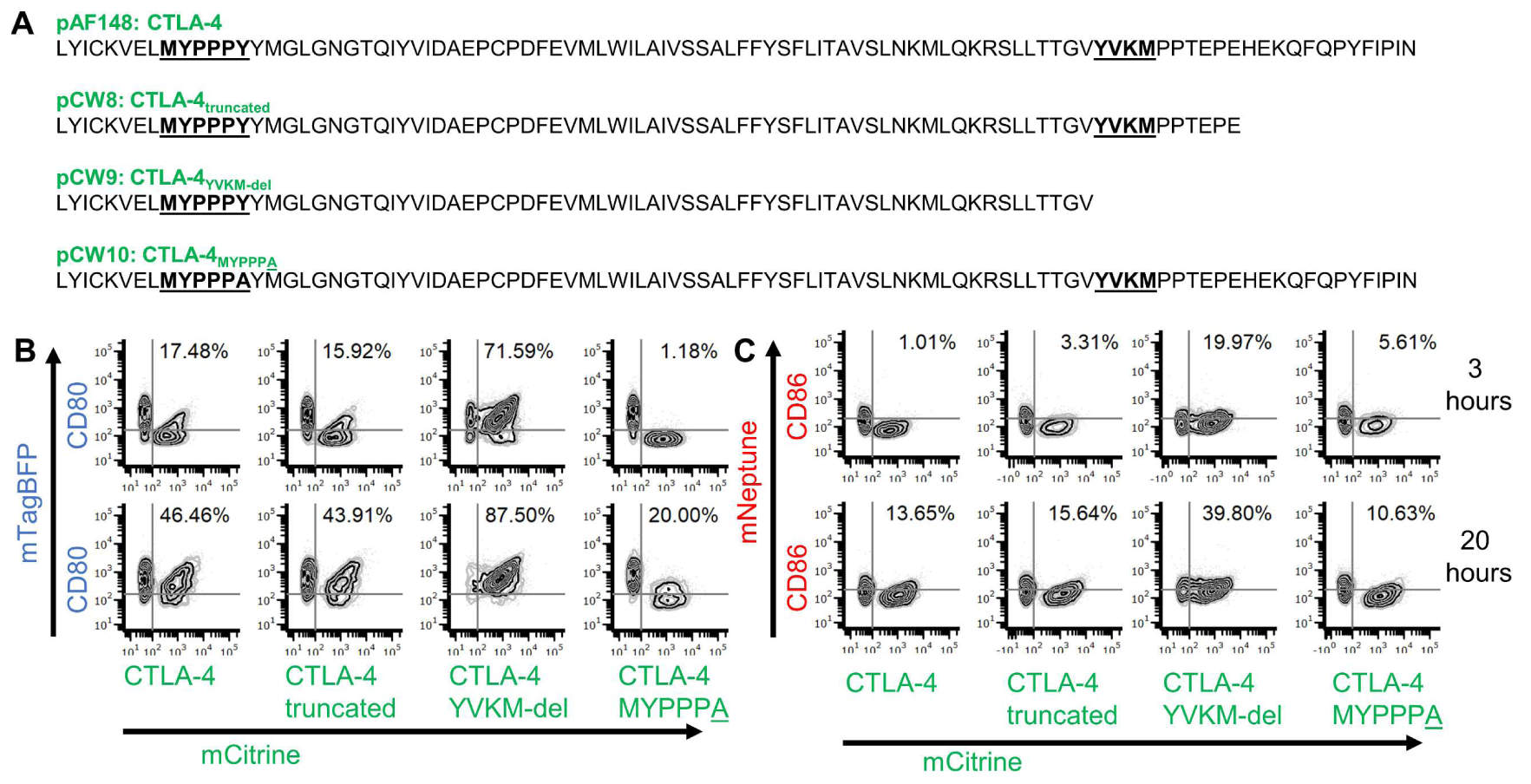
CTLA4 mutants show key binding and protein cycling motifs are conserved. (A) C-terminus sequence alignment of CTLA4 wildtype (WT) and variants (truncated, CTLA4_YVKM_ removal and CTLA4_MYPPP<u>A</u>_) used in this study. (B) Intercellular protein transfer (IPT) is observed in 3-hour and 20-hour co-culture assays of CTLA4 WT and CTLA4 truncated with CD80 (one-way); and CTLA4_YVKM-del_ and CD80 (bidirectional). No or minimal protein exchange is observed between CTLA4_MYPPPA_ and CD80. (C) No clear indication of IPT is observed between CTLA4 WT and CTLA4 truncated with CD86 in 3-hour assay, but minimal protein exchange is observed in 20-hour assay. Bidirectional protein transfer is observed between CTLA4_YVKM-del_ and CD86; whereas minimal protein transfer is shown in CTLA4_MYPPP<u>A</u>_ and CD86 co-culture assays. Results shown are representative of n=2 technical replicates.

In all species studied to date, the MYPPPY motif is required for ligand binding, so we hypothesized that IPT would be reduced in cocultures using the CTLA4_MYPPPA_ mutant compared to other CTLA4 variants (**Fig. 2A**). Coculture of CD80 with mutant CTLA4_MYPPPA_ showed minimal or no protein transfer (3-hours: 1%; 20-hours: 20%) (**Fig. 2B**), suggesting that the MYPPPY motif is critical for CTLA4-CD80 binding interactions in Tasmanian devils. The coculture of devil CTLA4 WT and CTLA4 mutants with CD86 ligand did not reveal a clear indication of intercellular protein exchange in the 3-hour assay even though the flow cytometry results showed a slight shift of CTLA4 (WT and mutants) to the UR quadrant (**Fig. 2C**). However, 20-hour coculture assays revealed a one-way protein exchange from CD86-mNeptune to CTLA4 WT-mCitrine (14%) and CTLA4 truncated-mCitrine (16%) (Fig. 2C). Additionally, cells expressing mutant CTLA4_YVKM-del_ cocultured with CD86 showed bidirectional protein transfer between both cell lines in both 3-hour (20%) and 20-hour (40%) assays (Fig. 2C). Lastly, CD86-mNeptune cocultured with mutant CTLA4 MYPPP<u>A</u> exhibited minimal protein exchange in 20-hour assay (11%) (**Fig. 2C**).

CTLA4 orthologues for 233 species are listed in the GenBank data base (accessed 8-May-2020; Table S3) (Benson et al., 2012). The CTLA4_MYPPPY_ and CTLA4_YVKM_ motifs are conserved in 137 and 138, respectively, of the 140 mammalian species. The CTLA4_MYPPPY_ motif is conserved in all 76 bird species in GenBank with CTLA4 orthologues, and the CTLA4_MYPPPY_ motif is conserved in 75 bird species. Only 3 reptiles have the CTLA4_MYPPPY_ motif and 4 have the CTLA4_YVKM_ motif. All three amphibians have the CTLA4_MYPPPY_ motif, but none have the CTLA4_YVKM_ motif. The coculture assays described here is amenable for functional testing of cell surface interactions in any of the above species, without the need to purify recombinant proteins. However, primary cells for each species or amphibian, reptilian, avian cell lines should be used instead of mammalian cell lines.

In summary, our results and analyses suggest that strong stabilizing selection has operated to maintain the function of these key immune checkpoint genes. This opens the door to immunotherapies that target these proteins for most mammalian species, and potentially for most bird species and some reptiles and amphibians. Although we have focused on marsupials in this manuscript, we believe that these methods can potentially be applied to the study of receptor-ligand interactions in invertebrates. We have published step-by-step protocols to facilitate comparative immunology studies in other non-model species (Flies et al., 2020b).

## Abbreviations

IPT: intercellular protein transfer
DFT: devil facial tumour
IFNγ: interferon gamma
MHC: major histocompatibility complex
CTLA4: Cytotoxic T-Lymphocyte Associated Protein 4
BFP: blue fluorescent protein

## Supplementary materials

**Table S1. Summary of genes and plasmids**

**Table S2. Primer sequences for plasmid assembly**

**Table S3. Primer sequences for Sanger sequencing**

## Acknowledgments

We Amanda Patchett, Camila Espejo, Chrissie Ong, Rob Gasperini, and Ruth Pye for assistance in the laboratory and general advice on this project.

## Author contributions

ASF and CW designed the study; ASF, CW, JMD, PRL, PRM, and TLP developed the technology; CW performed the experiments; ASF and CW created the figures; ASF, CW, TPL, ABL, and GMW analyzed the data; CW, ASF, and GMW wrote the manuscript and all authors edited the manuscript.

## Availability of data and materials

Data is available upon request, and will be made freely available in the University of Tasmania Research Data Portal (https://rdp.utas.edu.au) following peer-reviewed publication.

